# REPTRA: Mapping Immune T Cell Receptor Activity from Full Sequences with a Debiased Contrastive Loss

**DOI:** 10.1101/2025.10.22.683939

**Authors:** John Abel, Brinda Vijaykumar, Neel Patel, Anthony J. Coyle, Christophe Benoist, Daniel C. Pregibon

## Abstract

Contrastive learning is amenable to representing the vast space of interaction between highly specific T cell receptors (TCRs) and the epitopes to which they bind, potentially enabling diverse applications in immune engineering. However, progress in mapping TCR-epitope recognition has been limited by skewed datasets and training approaches that may exacerbate biases and obfuscate model performance. Furthermore, most TCR-epitope models represent only one of three complementarity-determining regions (CDRs) of the TCR, potentially limiting performance. Here, we present a CLIP-style contrastive-learning model for representation of epitopes and T cell receptor activity (REPTRA), incorporating full TCR sequences and trained using a debiased InfoNCE loss. We trained this model on a dataset with over fivefold more epitope diversity than previous reports, collected using the DECODE platform (Repertoire Immune Medicines). We demonstrate resulting improvements in model performance, and ablate the modified loss and TCR representation to demonstrate the contributions of this approach. Furthermore, we apply an interpretability analysis to the REPTRA attention-pooling projection heads to reveal that CDR1 and CDR2, in addition to CDR3, are important for learning TCR-epitope recognition. In doing so, we develop a performant model for contrastive mapping of T cell receptor activity.

## 1 Introduction

The immune synapse is the biological interface central to the body’s cellular reponse in cancer, autoimmune disorders, and infectious disease. In the immune synapse, a T cell receptor (TCR) recognizes a peptide from a degraded protein presented on a cell’s surface by a major histocompatibility complex (MHC) molecule [1]. TCRs that recognize and bind specific peptide-MHCs (pMHCs) initiate an immune response. Deciphering the immune synapse would enable engineering a variety of immune therapies, e.g. beneficial activation of adaptive immunity (for TCR-mimic therapeutics) or modulation of immune response to self-antigens (for immune-tolerizing vaccines) [18, 23, 33]. However, this remains a challenge due to the diversity of synapse features. To date, approximately 10^10^ TCRs and 10^6^ pMHCs have been described [36]. Machine learning approaches for deciphering the interaction of TCRs and pMHCs in the immune synapse from sequences, i.e. contrastive mapping, is therefore highly desirable.

Numerous recent studies have sought to directly predict TCR-epitope specificities with a variety of machine learning-based methods [4, 8, 10, 13, 15, 21], and several have made significant progress in tackling this challenge. However, data composition and problem structure have posed significant challenges for applying machine learning in this space. Despite the wealth of available TCR sequences, reported TCR-pMHC interactions are limited. Few well-characterized epitopes (peptides presented by MHC) from common viral antigens such as influenza, Epstein-Barr virus, cytomegalovirus, and SARS-CoV-2 comprise the majority of large public databases of positive TCR-epitope interactions [32, 35]. Generating negative TCR-epitope interactions is additionally challenging: for example, sampling negative TCRs from a healthy TCR repertoire leads to distribution shift and loss of generalization [21]. Contrastive learning suits the overall problem structure given the very low positive-to-negative interaction ratio, yet direct application of typical contrastive learning techniques where each positive TCR-epitope interaction is used to create a similarity matrix leads to a quadratic weighting of epitopes with respect to the number of occurrences of their corresponding TCRs. This results in a latent space biased toward the few highly studied antigens and potentially inflated performance metrics. If a model can accurately predict non-interactions for the few epitopes that comprise *>*50% of the test data, few-shot performance will appear strong, yet the model will fail in application.

In this study, we present several modifications to the TCR-epitope contrastive learning problem with a CLIP-style architecture. We applied our approach to a diverse TCR-pMHC dataset collected using the DECODE platform for studying TCR-pMHC interaction (Repertoire Immune Medicines) [12]. We contribute the following:

1. a contrastive model that leverages the entire TCR amino acid sequence with learned weighting via attention pooling,
2. epitope-frequency-debiased InfoNCE loss and evaluation metrics enabling improved contrastive learning of the TCR-epitope interaction space, and
3. empirical scaling and evaluation results on a larger TCR-epitope interaction dataset demonstrating the utility of these methods and identifying limitations.

Together, these methods enable training a performant model for contrastive representation of epitope and T cell receptor activity (REPTRA).

## 2 Related Work

Numerous recent studies have sought to directly predict TCR-epitope specificities from sequence or structure with a variety of machine learning-based methods and the ImmRep challenge is a recurring venue for the development models for predicting TCR-epitope specificity [22, 27].

Sequence-based methods have commonly been used for TCR-pMHC binding prediction. Croce *et al*. proposed MixTCRpred, a suite of transformer-based pMHC-specific binary classification models for predicting TCR binding to the pMHC of interest [4]. Meynard *et al*. proposed TULIP, a unique probabilistic method for predicting TCR-epitope-MHC interactions allowing the inclusion of incomplete data [21]. Importantly, this study revealed challenges in evaluating TCR-epitope models, and proposed several methods for alleviating evaluation biases. The NetTCR family of models has also made steady progress toward TCR-pMHC mapping using a variety of deep learning methods tailored to the TCR-pMHC space [25–27]. Finally, Im *et al*. proposed ImmuneCLIP, a CLIP-style contrastive learning model for TCR-epitope recognition [13]. ImmuneCLIP presents a strong foundation for contrastive learning of TCR-epitope interactions, showing that a CLIP-like formulation meets or exceeds performance of other methods while identifying key design parameters for this model. In other contexts protein-lanuage-model-based CLIP-style methods have shown utility in predicting peptide-protein interaction [29].

Structure-based prediction of TCR-pMHC binding has more recently come into focus. Recent successes in developing pMHC minibinders [16, 20] and advances in *de novo* antibody design [24, 34] lend optimism that TCR-pMHC specificity may be deciphered via structural prediction. To this end, recent studies have used folding models in conjunction with lightweight prediction using fold characteristics to predict TCR-pMHC binding [7]. Other studies have focused on developing improvements in TCR-pMHC docking to improve docking metrics and decrease computational cost vs. best-in-class general structural models such as AlphaFold-Multimer [11, 37]. However, the flexible and possibly dynamic nature of CDR-peptide interaction, sparsity of TCR-pMHC crystal structures, and relatively high computational cost of TCR-pMHC docking present challenges, especially for screening extreme volumes of potential interactions.

Broadly, a challenge in TCR-epitope binding prediction is how to perform negative sampling for training and evaluating TCR-epitope models [14, 21]. Previous approaches have included sampling from healthy TCR repertoires or creating negatives from other TCRs or epitopes in the dataset within a given split, due to extremely low likelihood that any given TCR will bind any one epitope [4, 13, 21, 27]. However, each of these methods has shown notable bias. Healthy-repertoire sampling has been shown to result in learning biases associated with the healthy TCR samples, rather than learning biological representations of TCR-epitope interaction [21]. Meanwhile, randomly sampling negatives from the test dataset leads to overrepresentation of common epitopes among the negative samples. In one instance, simply counting the occurrences of an epitope in the test set yielded a highly performant model [27], and another where at least one third of test dataset comparisons were TCRs vs. the common influenza epitope GILGFVFTL or another epitope vs. GILGFVFTL-reactive TCRs, which may inflate apparent performance on rare epitopes due to confident prediction of easy negatives. In these cases, reported performance on “unseen epitopes” may be due to correctly eliminating TCRs that bind common epitopes, rather than correctly identifying positive TCR specificities. Previous work has found that model performance depends largely on the imbalance within the train and test datasets of choice, with low generalization [9]. Overall, a lack of field-consensus benchmarks causes challenges in fair evaluation of proposed methods.

## 3 Approach

### 3.1 Datasets

We aggregated multiple datasets for model training and evaluation. We selected all TCR-epitope pairs from the VDJdb database with a confidence score of 3 [32]. To reconstruct TCR alpha and beta subunit sequences from a given V-gene, J-gene, third complementarity-determining region (CDR3), and C-gene in VDJdb, we aligned the CDR3 amino acid sequences with the reference V/J/C-gene amino acid sequences from the Immunogenetics (IMGT) database [2] via CellRanger (10X Genomics). To significantly expand upon commonly used public data, we identified TCR-epitope specificities using Repertoire Immune Medicine’s DECODE^TM^ platform yielding interactions between over 10M TCRs and thousands of epitopes as measured from patient samples across disease states in a manner consistent with [12]. Here, full TCR sequences were generated directly. To enable a contrastive framework, we selected only positive TCR-epitope pairs from these data with the highest confidence epitope recognition, representing approximately 0.0005% of all measured interactions. This ultimately yielded a dataset with approximately 7 × the epitope diversity of other sources.

**Table 1.** dataset composition for REPTRA training and evaluation, and comparison with the TULIP benchmark. Note that unique positive TCR-epitope pairs are reported, because negative sampling procedures may vary and duplicate positive pairs are included in the TULIP dataset. To demonstrate dataset skew, the number of TCRs pairing to the three most prevalent epitopes for the R-1 and TULIP datasets (respectively) are reported.

|  | **R-1** |  |  | **TULIP** |
| --- | --- | --- | --- | --- |
|  | Train | Val | Test | Test |
| Unique Positive Pairs (N) | 13,634 | 2,352 | 2,754 | 854 |
| TCRs (N) | 12,254 | 2,101 | 2,451 | 853 |
| Peptides (N) | 4,416 | 767 | 793 | 148 |
| KLGGALQAK (N, %) | 1407 (10.3%) | 305 (13.0%) | 300 (10.9%) | 0 (0.0%) |
| GILGFVFTL (N, %) | 1036 (7.6%) | 218 (9.3%) | 420 (15.3%) | 310 (36.3%) |
| RAKFKQLL (N, %) | 316 (2.3%) | 70 (3.0%) | 80 (2.9%) | 0 (0.0%) |
| NLVPMVATV (N, %) | 121 (0.9%) | 26 (1.1%) | 47 (1.7%) | 68 (8.0%) |
| GLCTLVAML (N, %) | 271 (2.0%) | 66 (2.8%) | 75 (2.7%) | 66 (7.7%) |

We split the resulting dataset (“R-1”) by first holding all TCRs in the TULIP test dataset out for testing. We assigned TCRs pairing with epitopes that had two or fewer TCRs to the training dataset, for maximal training diversity. We split all remaining unique TCRs for each epitope into training (70%), validation (15%) and held-out test (15%). Following data splitting, we confirmed that no TCRs occurred in multiple dataset splits, and that each TCR-epitope pair was unique. R-1 dataset composition was found to be diverse and less dependent on any individual epitope (Tab. 1). As is common in contrastive learning, we treated all non-positive TCR-epitope interactions as implicitly negative pairs. We did not provide MHC information to our model or use the MHC in constructing the loss, treating the peptide as the sole representative of the pMHC epitope. Less than 6.5% of peptides were presented by more than one MHC molecule; we therefore speculated that adding MHC identity would not be required for pMHC recognition (as in [13]).

### 3.2 Model Architecture

We constructed a simple contrastive learning model for mapping TCR-epitope reactivity (shown in Fig. 1A). Briefly, an epitope is encoded by passing the peptide sequence through an ESM-2 encoder followed by an attention pooling and a linear projection into the latent space [19]. A TCR is encoded by passing the amino acid sequence for the alpha and beta subunits through respective ESM-2 encoders, followed by attention pooling, concatenation, and projection into the latent space. The third complementarity-determining region (CDR3) of each TCR subunit is thought to be the primary (but not sole) driver of TCR-epitope binding [3], thus, we trained models that encoded the CDR3 alone as well as the full TCR sequence.

**Figure 1.**
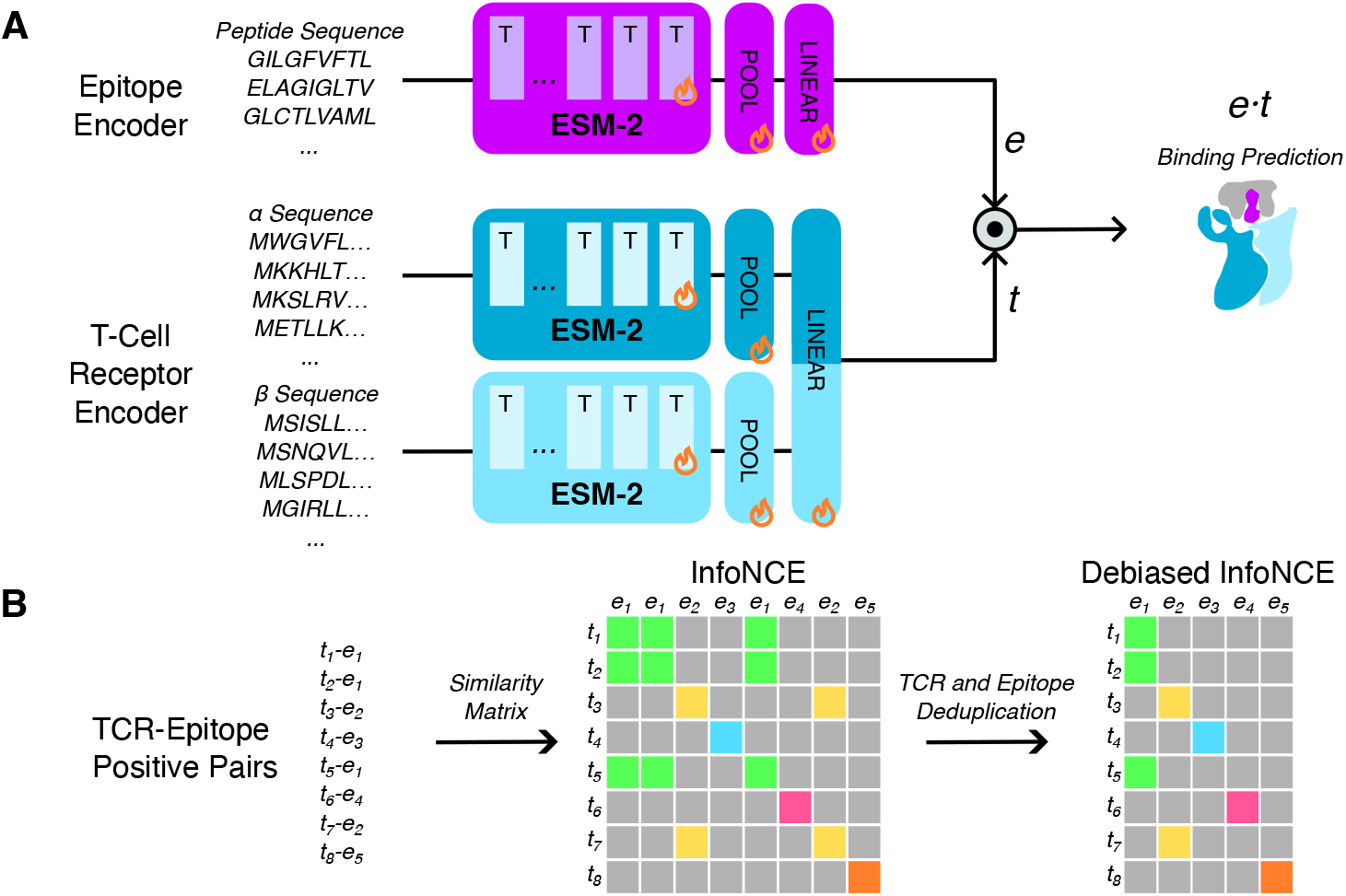
(A) schematic of REPTRA structure. Flames denote which blocks are trained. Note that the MHC is not explicitly encoded. (B) Visualization of debiased InfoNCE loss. By deduplicating epitopes and TCRs, we eliminate duplicate entries in the contrastive loss and evaluation.

### 3.3 Training and Implementation Details

All models were developed in PyTorch [31] and trained in a consistent manner using the AdamW optimizer [17] with hyperparameters provided in Tab. A1. An epitope-frequency-debiased InfoNCE loss was defined and used for model fitting, as detailed in Section A.3. We selected the best model for each training run using the debiased InfoNCE AUROC on the validation split. We defined the total number of tokens used per training run as the sum of the epitope and TCR tokens for each TCR-epitope pair passed through the model. All models were trained on a single NVIDIA A10 GPU using mixed precision; a training run was complete in approximately 5 hr for a full-TCR model or 1 hr for a CDR3-only model.

### 3.4 Code Availability

Code for training and deploying REPTRA-style models is available under a non-commercial license at https://github.com/repertoireimmunemedicines/reptra. TCR-epitope affinity data will not be shared due to patient privacy and IP considerations.

## 4 Experiments and Results

### 4.1 Scaling TCR-Epitope Contrastive Learning

We tested whether TCR-epitope contrastive learning was limited by data or by encoder model capacity by training REPTRA architectures using ESM-2 backbones from 8M to 150M parameters (Tab. A2). We compared validation loss and debiased AUROC between these backbones to identify potential scaling laws for TCR-epitope contrastive learning. Surprisingly, we found that increased backbone size and model capacity did not improve validation performance when training on CDR3 regions alone or on full TCR sequences (Fig. A1), indicating that even with a much larger dataset than previous studies and richer description of the TCR, we are limited by the diversity of the data. Furthermore, we observed the encoding full TCR sequences dramatically improved validation loss and AUROC. We selected the full-TCR REPTRA model with ESM-2 150M (“REPTRA”) and the CDR3 REPTRA model with ESM-2 150M (“REPTRA-CDR3”) for further evaluation due to slightly smoother training and for consistency between CDR3 and full-TCR models.

**Table 2.** evaluation metrics for REPTRA, TULIP [21], and ImmuneCLIP [13] on the TULIP and R-1 test datasets. All values are macro AUROCs. We report performance on the TULIP test dataset as a benchmark, however 568/853 (66%) of TULIP test TCRs were also in the ImmuneCLIP train split (see Section A.2). Thus, there is significant leakage inflating ImmuneCLIP performance. Note that REPTRA cannot be applied to the TULIP test dataset, which lacks non-CDR3 information.

| Model | TULIP Test | R-1 Test |  |
| --- | --- | --- | --- |
|  | T-AUROC | I-AUROC | D-AUROC |
| REPTRA | - | **0.967** | **0.886** |
| REPTRA-CDR3 | 0.805 | 0.942 | 0.820 |
| TULIP | 0.732 | 0.887 | 0.762 |
| ImmuneCLIP | 0.804 | 0.751 | 0.676 |

### 4.2 Model Evaluation and Benchmarking

We evaluated REPTRA alongside to two top-performing models in the literature on the publicly available TULIP test dataset and our R-1 test dataset (Tab. 2). Benchmarking is a known challenge for ML models of TCR-epitope recognition, as each model has been trained on TCRs paired with different epitopes, and there is known train-test leakage. We additionally used our full R-1 test dataset as a challenge to probe performance across a wider portion of epitope-space. We calculated AUROC for TULIP test using the exact TULIP negatives (T-AUROC), and for R-1 test using both the confusion matrix as in the InfoNCE loss (I-AUROC) and deduplicated as in the debiased InfoNCE loss (D-AUROC). As in the scaling experiments, REPTRA outperforms REPTRA-CDR3 indicating that non-CDR3 regions contribute to binding prediction. REPTRA and REPTRA-CDR3 outperform comparator models in the literature, even outperforming TULIP on the TULIP test dataset. The less-performant REPTRA-CDR3 (with full representation ablated) still performs comparably to ImmuneCLIP on the TULIP test dataset, even though 568/853 (66%) of TULIP test TCRs appear in the ImmuneCLIP training dataset due to leakage in data splitting (see Section A.2 for a longer discussion).

### 4.3 Ablation Experiment

We performed an ablation study to better understand the contributions of debiased InfoNCE loss and full-TCR representation, reporting validation D/I-Loss and D/I-AUROC, and test D/I-AUROC. We ablated the debiased InfoNCE loss (A1, replacing it with standard InfoNCE and using InfoNCE loss as the model selection criterion), and both debiased InfoNCE and the full-TCR representation (A2, replacing it with the typical CDR3-alone encoding). Results in Tab. 3 indicate that each of these adjustments to the model contribute to its overall performance. Interestingly, using the debiased loss improves model performance on the standard InfoNCE-style AUROC, supporting the notion that including duplicated comparisons distorts the learned latent space.

**Table 3.** ablations of REPTRA. We replaced debiased InfoNCE with standard InfoNCE and the full-TCR sequence with only CDR3s, finding that training on standard InfoNCE loss or CDR3s alone results in diminished test-time performance.

| Model (Ablation) | R-1 Validation |  |  |  | R-1 Test |  |
| --- | --- | --- | --- | --- | --- | --- |
|  | I-Loss | I-AUROC | D-Loss | D-AUROC | I-AUROC | D-AUROC |
| REPTRA (None) | 3.65 | **0.963** | **5.59** | **0.867** | **0.967** | **0.886** |
| A1 (InfoNCE) | **3.54** | 0.933 | 5.84 | 0.844 | 0.946 | 0.864 |
| A2 (InfoNCE/CDR3) | 3.89 | 0.857 | 6.31 | 0.762 | 0.896 | 0.787 |

### 4.4 Interpreting REPTRA Attention

We initialized the attention pooling weights within REPTRA to be a mean pool. During training, the model learns to attend to the regions of the sequence useful for determining epitope specificity. Because non-CDR3 information from the full TCR sequence was found to contribute to model performance, it is of interest to determine how REPTRA attends to the full TCR sequences. We tested this by extracting the weights from attention pooling on TCRs within the test dataset (Fig. 2). We found that in addition to CDR3s, TCR sequence regions consistent with CDR1/CDR2 positions are weighted higher during attention pooling than the framework regions throughout, especially for the TCR beta chain. We investigated further by structurally modeling a GILGFVFTL-binding TCR and mapping the attention weights for that TCR onto its structure (shown in Fig. 2B) [6, 30]. We found that the highest-attention regions were those oriented toward the pMHC complex. When reviewing the GILGFVFTL-binding TCR, we found an additional high-attention region corresponding to the sequence REKKESFPL within the Framework-3 region of the beta chain (IMGT positions 80-87), which has previously been referred to as “CDR2.5,” and may contact the epitope [5].

**Figure 2.**
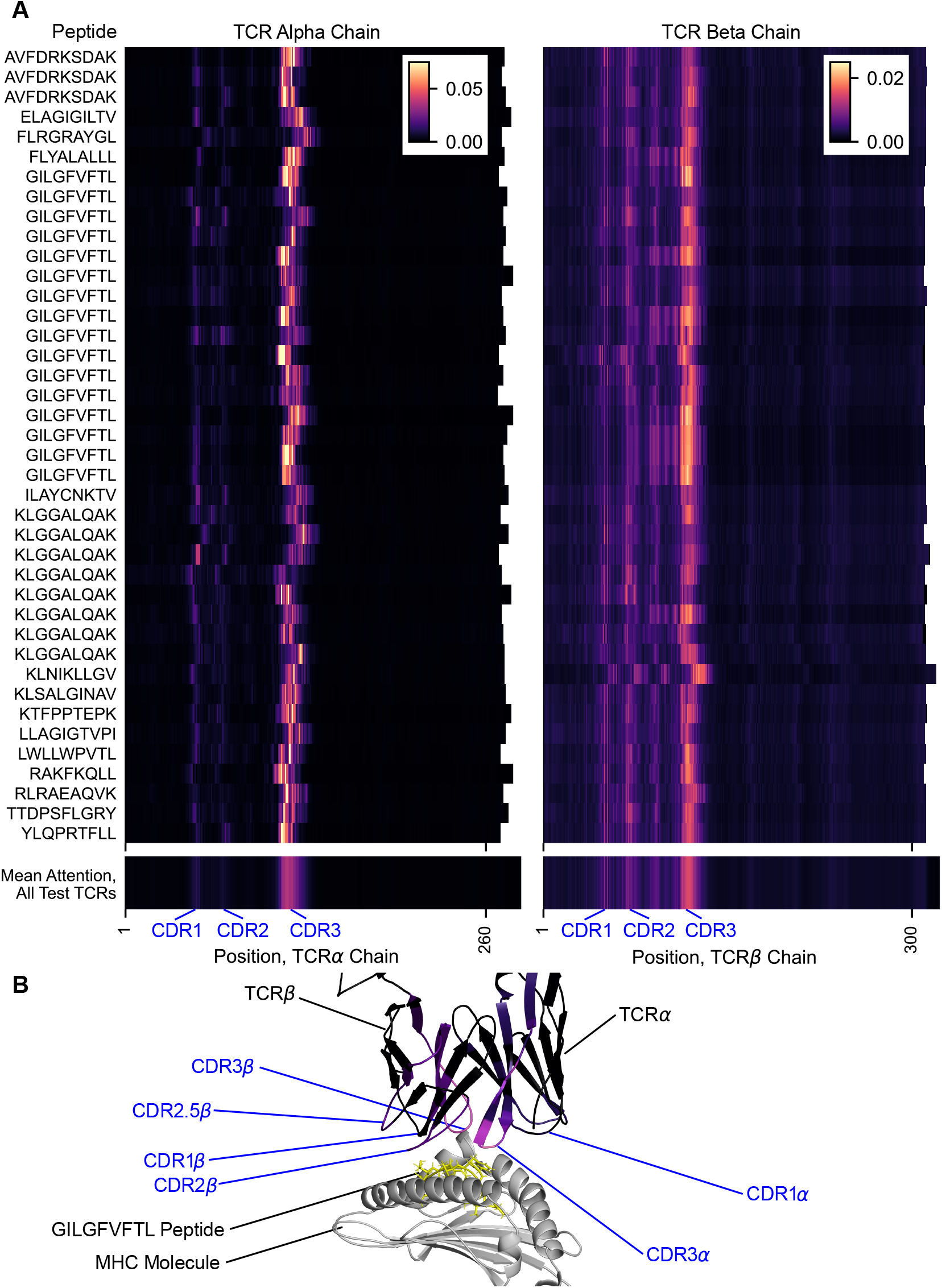
(A) a heatmap of weights across the TCR alpha and beta chains for 40 randomly selected TCRs mapping to varied epitopes. Mean weights on each position in the TCR alpha and beta chains are shown below, computed across the entire R-1 test dataset. Note that we include the TCR leader peptide in our full TCR sequence, although this region appears to receive very low attention. The CDR3 positions may vary between TCRs due to differences in the leader/variable region length. (B) Attention mapped onto structural model of TCR-pMHC for an example GILGFVFTL-binding TCR.

### 4.5 Limitations and Future Work

A limitation in this study (and to-date all other studies of TCR-epitope recognition) is sparse coverage of the epitope space, as indicated by lack of performance improvement with larger encoder backbones. Although our R-1 dataset includes approximately 7 × the diversity of literature examples, this represent a small fraction of known epitope space. Furthermore, our model does not incorporate MHC information. Because most peptides are only presented by one MHC, we did not add this representation to our initial model.

Importantly our approach relies on sequence alone, and although ESM-2 encodes structural information from sequences, no direct structural metrics are used. We made this decision due to well-known difficulty in correctly identifying CDR loop folding (which may take many, dynamic conformations) and the significant computational effort required to dock up to millions of TCRs with hundreds of thousands of peptides with thousands of MHCs. Future approaches might leverage a two-stage process, where a high-throughput contrastive learning model maps plausible TCR-pMHC affinities, and a refined structural model tuned to TCR-pMHC docking enables precise screening of the candidate TCR-pMHC pairs.

## 5 Conclusion

We have presented two modifications to the TCR-pMHC contrastive learning problem: representation of the full TCR sequence and a modified contrastive loss function. Our approach yields improved model performance and enables interpretation of model weights in a way that is consistent with known biology. Furthermore, it indicates that existing datasets lacking information outside the CDR3 may not be optimal for developing TCR-epitope ML models. It would be of interest to apply interpretability approaches to answer specific scientific questions, e.g. which epitope residues are recognized by a TCR (to thus identify undesirable cross-reactivity), or determine whether one or more CDR motifs is associated with recognizing a given epitope. Future work in this space may benefit most significantly from a more diverse set of TCR-pMHC training data, broadly covering self-antigens presented in Class I and Class II MHCs in addition to the limited set of well-understood viral epitopes, with the ultimate goal of enabling broad generalization beyond known epitopes. To this end, we plan to scale by leveraging a larger fraction of our TCR-pMHC interaction database toward developing a broadly generalizable model.

## Acknowledgments

We gratefully thank the Lab Platform team at Repertoire Immune Medicines (Amy Perea, Genesis Tejada, Del Leistritz-Edwards, William Ruff, Qiaomu Tian, Jack Prazich, Veronica Sanchez, Katherine Triebel, Marcos Wille, Manning DelCogliano, and Preet Joshi) for data acquisition and biological direction. We additionally thank the Computational Engineering team (Kane Hadley, Elysha Sameth, Aswathy Sheena, and George Regas) for ML Ops and data support.

## Disclosures and Funding

JA, BV, NP, AC, CB, and DC are employed by and/or have received equity in Repertoire Immune Medicines, Inc.

## A Appendices and Supplementary Material

### A.1 Scaling

**Figure A1:**
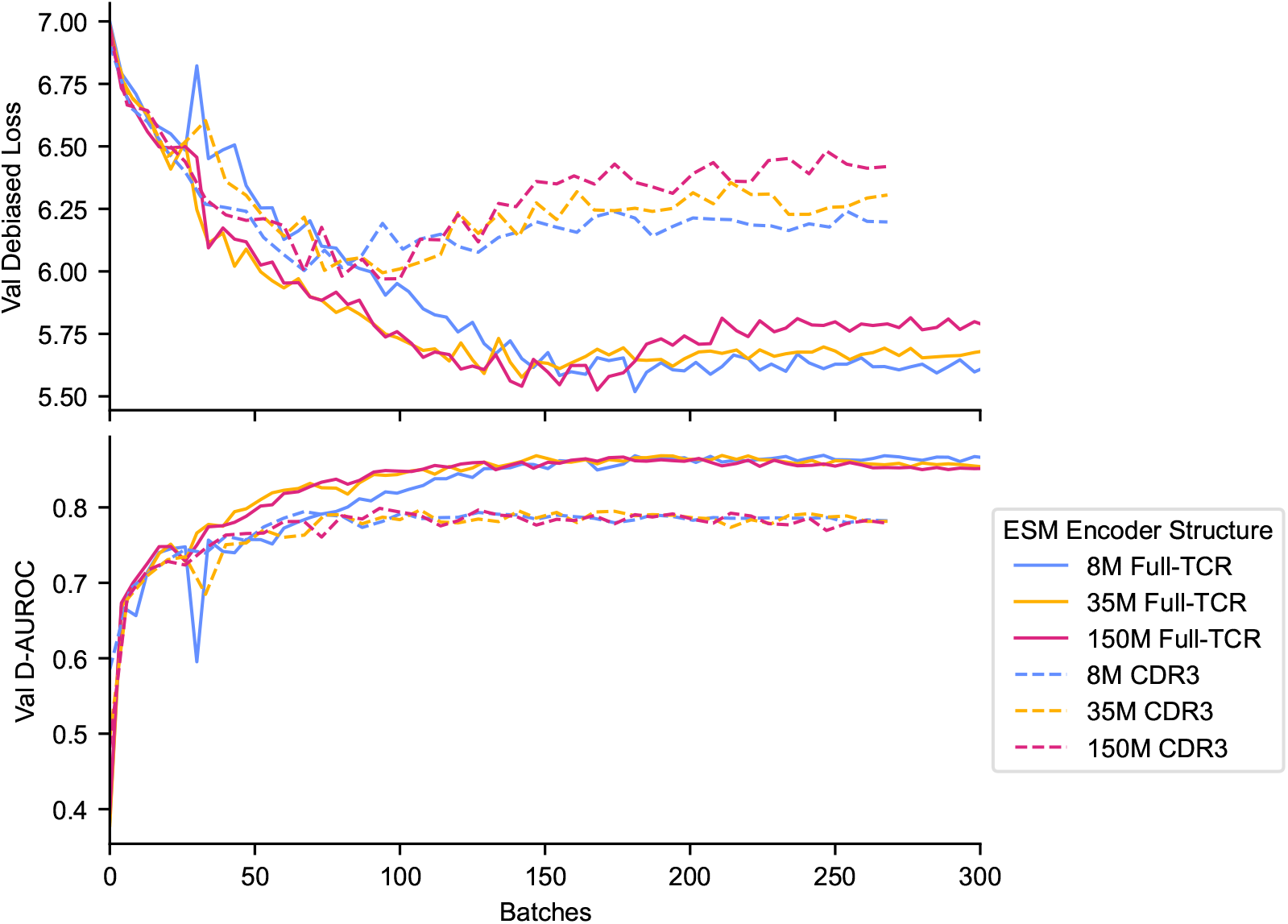
scaling REPTRA encoders. We show performance vs. batch (optimizer step) because tokens per batch varies approximately 10-fold depending on whether the full TCR or CDR3-only is used.

#### A.2 Data Splitting and Leakage

Prior studies have struggled to maintain consistent data splits in this space, in addition to the challenge of proper negative sampling. Here, we chose to treat the TULIP test dataset [21] as a true benchmark, rather than attempt to re-split such that all epitopes were seen by each model (as in [13] and elsewhere), in order to stay true to the spirit of a benchmark, i.e. so that all models can be evaluated on the exact same data.

This approach leads to one additional challenge: potential data leakage for models that are not explicitly split consistently with TULIP in mind (i.e. ImmuneCLIP). ImmuneCLIP previously addressed this challenge by removing “all epitope/TCR pairs in each of the test sets that [the] model had seen during training.” Implicitly, ImmuneCLIP retained TCRs that were seen during training but were paired with different epitopes; thus it has been trained on some of the TCRs in negative TCR-epitope pairs in TULIP. We found that 568 of 853 TULIP test TCRs were in the ImmuneCLIP train split. Even if paired with different negative epitopes in the TULIP test dataset, the ImmuneCLIP model has seen that these TCRs are positive for other epitopes during training; thus there is significant information leakage that lends a positive bias to ImmuneCLIP. We nevertheless report the ImmuneCLIP performance here, as a conservative upper bound on the method.

#### A.3 Debiased InfoNCE Loss

A standard approach in contrastive learning is to use the InfoNCE loss [28]. Here, for each batch of *N* positive pairs, a similarity matrix is constructed of dimension *N* × *N* (where each anchor forms a row and each candidate forms a column). Cross-entropy is applied with the diagonal element typically considered the correct class. In bimodal settings, this is done in both directions and the losses are averaged. Equivalently, during evaluation, the *N* × *N* matrix is constructed and used to compute performance metrics (e.g. AUROC). It is established that many unique TCRs bind to the same epitope. Previous studies have addressed this by incorporating off-diagonal positives in the confusion matrix, up/downsampling, and maintaining a low batch size during training [13]. However, contrastive learning generally benefits from large batch sizes to learn the representation space, so keeping additional TCR diversity is helpful even for common epitopes. We further speculated that training using an InfoNCE-style similarity matrix would lead to distortion of the representation space due to duplicated comparisons with the same epitope.

If an epitope *e*_*a*_ forms positive pairs with *k* TCRs, in the InfoNCE confusion matrix, the number of positive and negative entries for *e*_*a*_ are given by:

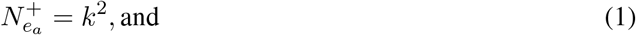

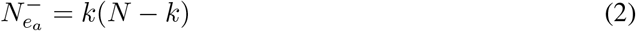

where the *k* unique positive TCRs and *N* − *k* unique negative TCRs are compared to epitope *e*_*a*_ a total of *k* times. During training common epitopes will dominate the representation due to nonlinear scaling with *k* (visualized in Fig. 1B) and during evaluation, the same negatives and positives may be over-counted in computing evaluation loss or other metrics. This results in apparently strong model performance even for TCRs that bind rare epitopes, but may be attributable to multiple comparisons with the same common negative epitope.

We developed a debiased InfoNCE loss by simply deduplicating epitopes and TCRs when constructing the InfoNCE-style confusion matrix. Using this approach, the number of positive and negative entries for *e*_*a*_ are given by:

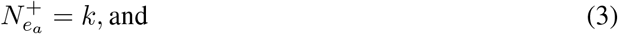

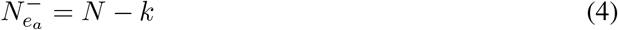

respectively. This enables larger batch sizes during training and ensures non-duplication of exact TCR-epitope comparisons during evaluation.

### A.4 Training Hyperparameters

For all model training, hyperparameters were held fixed with the exception of the maximal number of training tokens per run (and the evaluation interval), as noted, due to the tokens-per-sequence varying approximately tenfold between CDR3-only and full-TCR representations.

**Table A1:**
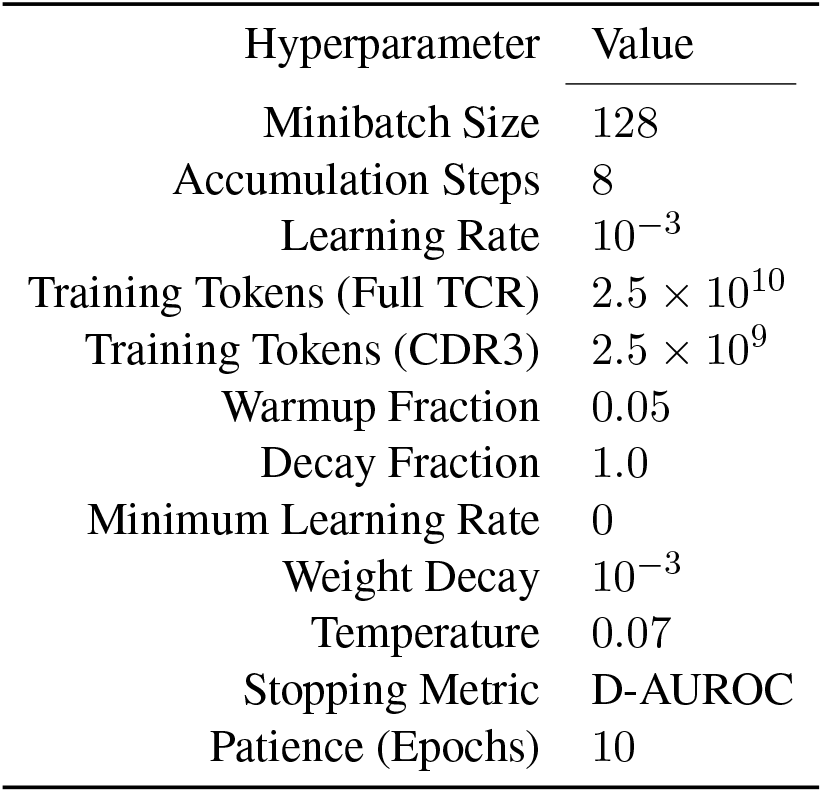
training hyperparameter values for training REPTRA.

| Hyperparameter | Value |
| --- | --- |
| Minibatch Size | 128 |
| Accumulation Steps | 8 |
| Learning Rate | $10^{-3}$ |
| Training Tokens (Full TCR) | $2.5 \times 10^{10}$ |
| Training Tokens (CDR3) | $2.5 \times 10^9$ |
| Warmup Fraction | 0.05 |
| Decay Fraction | 1.0 |
| Minimum Learning Rate | 0 |
| Weight Decay | $10^{-3}$ |
| Temperature | 0.07 |
| Stopping Metric | D-AUROC |
| Patience (Epochs) | 10 |

### A.5 Model Architectures

**Table A2:** model architectures tested for scaling REPTRA. Only the final transformer is fine-tuned.

| Model | ESM-2<br>Backbone | Projection<br>Head | Embedding<br>Dimension | Trainable<br>Parameters | Total<br>Parameters |
| --- | --- | --- | --- | --- | --- |
| REPTRA | 8M | Attention Pooling | 256 | 3.9M | 22.8M |
|  | 35M | Attention Pooling | 256 | 8.7M | 100.9M |
|  | 150M | Attention Pooling | 256 | 15.3M | 444.9M |

## Notes

https://github.com/repertoireimmunemedicines/reptra

